# The longer transmembrane helices of class I viral fusion proteins may facilitate viral fusion

**DOI:** 10.1101/2025.09.25.678357

**Authors:** Sahil Lall, Padmanabhan Balaram, M.K. Mathew, Shachi Gosavi

## Abstract

The viral envelope-resident viral fusion proteins (VFPs) fuse the envelope with the host cell membrane, which allows virus entry into host cells. The homotrimeric class I VFPs (cI-VFPs) are anchored to the envelope by three single-pass transmembrane helices (spTMHs). It is generally accepted that the hydrophobic length of a TMH matches the thickness of the hydrophobic membrane core, and this reduces TMH dynamics by stabilizing both the TMHs and the membrane. However, to enable fusion, a cI-VFP undergoes a large conformational transition which includes the disassembly of the spTMHs from a pre-fusion trimer and their subsequent reassembly with the cI-VFP fusion peptides into a post-fusion complex. Given the potential functional relevance of the cI-VFP spTMH disassembly-reassembly dynamics, we hypothesized that these spTMH lengths may be mismatched with the thickness of biological membranes. Here, we examine the predicted lengths of a curated dataset of cI-VFP spTMHs and find that they are indeed about 10% longer than the hydrophobic thickness of model membranes. The cI-VFP spTMHs are on average also longer than the spTMHs of human fusion and human non-fusion proteins. Since our dataset contains cI-VFPs from diverse enveloped viruses, and viral proteins are fast evolving, we conclude that the longer spTMHs are functionally relevant and thus, preserved. Our observation that the spTMH lengths of retrovirus-derived human fusogenic proteins, syncytins, are similar to those of cI-VFPs, supports this conclusion. The longer spTMHs may facilitate the cI-VFP conformational change and perturb the fusing membrane, reducing the barrier to fusion.

## INTRODUCTION

Membrane fusion is a complex process by which two separate biological membranes merge into one continuous membrane (*1–4*). Fusion is enabled by a complex of proteins constituting the “fusion machinery” which serve to dock the two membranes and bring them close enough for fusion to occur (*5–7*). The fusion machinery can further facilitate fusion by perturbing the structure of the fusing membranes (*2, 7, 8*).

In enveloped viruses, the viral membrane (known as envelope) fuses with a cellular membrane (e.g. plasma membrane, endosomal membrane, etc.) to release the viral genome into a host cell and initiate viral infection. The presence of only a single type of protein, the viral fusion protein (VFP), on the virus surface is sufficient to achieve viral fusion (*4, 9*). VFPs are large proteins that enable membrane fusion by binding host cell receptors and bringing the viral envelope close to the host cell membrane. Based on their structure, stoichiometry, and mechanism of action, VFPs are classified into different classes (*4, 6, 10, 11*). The VFPs of several well-studied viruses such as hemagglutinin (HA) of influenza (*12*), Env/gp160 of HIV (Human Immunodeficiency Virus) (*13*), and the spike protein of Coronaviruses (CoVs) (*14*) are class I VFPs (cI-VFPs).

The cI-VFPs are homotrimeric. In their pre-fusion state (Figure 1A) each protomer of the trimer is anchored to the viral envelope through one single-pass transmembrane helix (spTMH) located near the C-terminus of the cI-VFP. Together, these three spTMHs are known as the transmembrane domain (TMD). In the pre-fusion state of the cI-VFP, large ectomembrane domains, N-terminal to the TMD bind the host membrane-resident receptor. This binding triggers a conformational change in the cI-VFP, exposing a fusion peptide within the N-terminal domains, which then inserts into the host membrane to initiate the process of viral fusion (Figure 1B). This extended conformation of the cI-VFP (Figure 1B) undergoes a conformational transition in which the ectomembrane domains fold back upon themselves leading to the formation of a six-helix bundle (Figure 1C) bringing the two membranes in apposition. The cI-VFP then transitions into its post-fusion state as the membranes fuse (Figure 1D).

**Figure 1.**
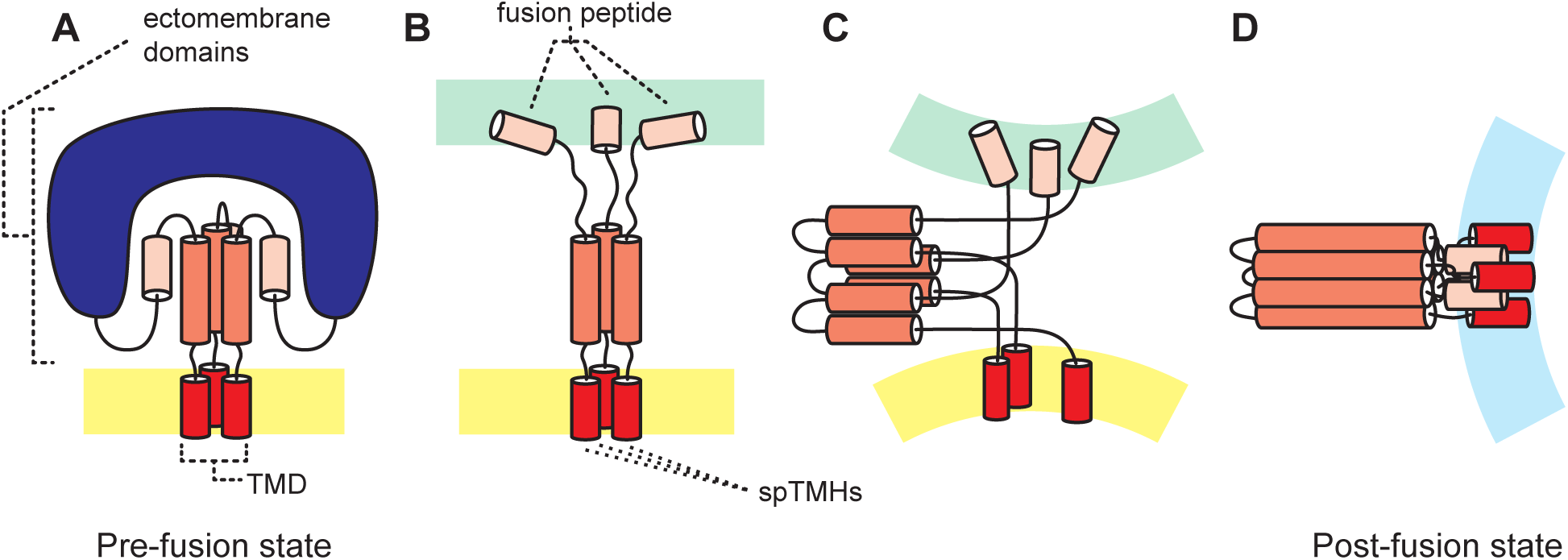
A simplified representation of viral fusion through a cI-VFP. (**A**) The pre-fusion state of a cI-VFP is its resting conformation on the virus surface. The pre-fusion state comprises a host cell receptor binding subunit (dark blue), and the fusion subunit which contains from N- to C-terminus: the fusion peptide (beige), the central stalk (orange) and the TMD (red). The TMD anchors the cI-VFP to the viral envelope (yellow in all the panels). (**B**) A lowered pH, receptor binding, and/or protease cleavage removes the receptor binding domain, freeing the fusion peptide (beige) to extend and dock with the host membrane (green in all the panels). (**C**) The extended conformation from (B) folds back on itself through the formation of a six-helix bundle (orange, left) that potentially perturbs the TMDs, deforms the viral membrane, and brings the virus and the host cell membranes closer together. (**D**) The post-fusion conformation of the cI-VFP is anchored to the fused membrane (light blue). The fusion peptides (beige) and the TMHs (red) interact in this state. For no single virus, are both the full-length pre-fusion (A) and post-fusion (D) states resolved.

The N-terminal cI-VFP ectomembrane domains undergo multiple large conformational changes (Figure 1) and have been structurally characterized in both their pre-fusion and post-fusion states in some viruses (*15, 16*). The dynamics of such ectomembrane domains have also been studied (*12, 17, 18*). However, both pre and post-fusion TMD structures are not available for any one cI-VFP (e.g. while the post-fusion structure of the well-studied wild-type SARS-CoV-2 spike with its TMD exists, only the isolated TMD with several mutations has been resolved in presumably the pre-fusion state). A complete characterization of the conformational conversion of the TMD from the pre-to the post-fusion state is also not available. Therefore, the process of bringing the two membranes in proximity through the dynamics of the cI-VFP ectomembrane domains is understood in many viruses but the mechanism of actual membrane fusion and the role of the spTMHs in facilitating fusion remains unclear. We recently found that the SARS-CoV-2 spike spTMH is unusually long, resulting in a hydrophobic mismatch with the thickness of a model bilayer membrane (*19*). Molecular dynamics simulations showed that this mismatch leads to spTMH dynamics (*19*). We hypothesized that these dynamics may promote fusion by both facilitating the spike’s conformational transition to its post-fusion state and perturbing the membrane. Further, if such hydrophobic mismatch-dependent dynamics are important for fusion, then the spTMHs across cI-VFPs should be long in order to preserve function. Here, we curate a dataset of cI-VFPs and analyze their predicted spTMH lengths.

Processes within eukaryotic cells such as fertilization, placental development, intracellular vesicular fusion and neurotransmitter release (*5, 9*) also require membrane fusion, and consequently, fusion machinery. But in contrast to the “one protein does it all” cI-VFPs, the cellular fusion machinery often consists of multiple proteins recruited to the membrane by the membrane-bound SNARE proteins (soluble NSF (N-ethylmaleimide sensitive factor) attachment protein receptors) (*7, 20, 21*). Similar to the cI-VFP protomers, many SNAREs are also anchored to the membrane through a single spTMH, with this helix helping the SNARE proteins get transported to and localized in the correct membrane (*22*). Based on their localization, different SNAREs are classified as t-SNAREs (localized to the target membrane) or v-SNAREs (localized to intracellular vesicles) (*21*). We compare the predicted spTMH lengths of human fusion proteins, such as human SNAREs to those of cI-VFP spTMHs.

More generally, bitopic or single pass transmembrane proteins (spTMPs) are proteins which can have large ectomembrane domains, but which have a single, helical membrane-spanning segment, the spTMH. Genome analyses have estimated that 15% of bacterial membrane proteins and up to 40% of human membrane proteins are spTMPs (*23*). The spTMHs are challenging to study and, as in cI-VFPs, much of the understanding of spTMPs comes from studies of their ectomembrane domains (*24*). However, it has become increasingly clear that not only the ectomembrane domains but also the spTMHs contribute to the function of the spTMPs. For example, the spTMHs are involved in the accurate membrane targeting of the spTMP (*25*), in protein-protein interactions within the membrane (*26, 27*), in the regulation of the activity of other membrane proteins (*28*), and in the formation of ion channels (*29*). Some of this spTMH function, such as breaking and forming protein-protein interactions, likely requires spTMH dynamics (*30, 31*).

Here, we first curate datasets of (a) cI-VFPs, (b) human fusion spTMPs and (c) human non-fusion spTMPs. Considering the potentially similar functional roles of the spTMHs of cI-VFPs and of human fusion proteins, and the diverse roles of human non-fusion spTMPs, we compare and contrast the spTMH lengths across the three datasets. Specifically, hydrophobic matching of the TMH length with the thickness of the membrane hydrophobic core is a strong signal for a polypeptide to be localized in that membrane (*32–35*). TMHs with a mismatched, long hydrophobic length localized in a membrane would have to adapt by tilting, segregating together, bobbing perpendicular to the membrane plane, etc. (*33, 36, 37*), and/or by rearranging nearby lipids (*34*). Dynamics created by the mismatched helix affects lipid ordering (*38*) and such a mismatch and the dynamics of cI-VFP TMDs (spTMH trimers) can disturb the membrane (*39*), which in turn can promote membrane fusion (*40*).

Both cI-VFP and human fusion spTMP function could be facilitated by a spTMH length-membrane thickness mismatch and the resulting spTMH dynamics. So, we first compared the average predicted spTMH lengths from multiple TMH prediction algorithms across these two datasets and found opposite effects. The predicted lengths of human fusion spTMHs are a good fit to their native biological membranes. Whereas, on an average, cI-VFP spTMHs are 2.3 residues longer than human fusion spTMHs, and presumably have a hydrophobic mismatch to their native membranes. Given the increasing evidence that both cI-VFP and human fusion spTMHs may locally disturb the membrane in order to enable fusion (*41*), we discuss the difference in average predicted lengths of the two datasets in the context of possible differences in mechanisms of membrane perturbation. The functions of human non-fusion spTMPs require a combination of spTMH anchoring/trafficking (which requires a fit to the membrane) (*35*) and dynamics (*30, 31, 42*), and thus, as expected, their predicted lengths are in-between those of the cI-VFP spTMHs and the human fusion protein spTMHs.

Overall, we find that the predicted lengths of cI-VFP spTMHs are longer than those of human spTMHs. These long spTMHs may both improve the cI-VFP conformational transition (*4, 5*) as well as locally destabilize the viral envelope (*19*) to facilitate the fusion of the viral envelope with the host membrane.

## METHODS

### Assembling curated datasets of single-pass transmembrane proteins (spTMPs)

We curated three spTMP sequence datasets. The first was a set of 65 cI-VFPs, the second set comprised 18 human fusion proteins that facilitate fusion in/of human cells, and the third “control” set contains 65 human spTMPs which have no known role in fusion and which are present on diverse eukaryotic membranes. The combined dataset contains 148 spTMPs and is given in the Supplementary Information. Unless otherwise noted, all sequence alignments were performed on the Clustal omega webserver hosted by EMBL-EBI (*43*).

#### cI-VFPs

The viral fusion protein sequences were obtained from the Uniprot database (*44*) using the query: class I viral fusion proteins (on 8^th^ August, 2024). The output of 886 reviewed sequences was filtered by existence at the “protein level”, which decreased the dataset to 351 sequences. Examination of the 351 protein sequences revealed that they contained several class II viral fusion proteins, and also proteins from organisms other than viruses that contained “class” or “virus” in their description, such as MHC class II transactivator, and endogenous retrovirus proteins. Therefore, using the 351 sequence set, 168 cI-VFP sequences were manually extracted. Incomplete sequences were removed from this set by filtering with the word ‘fragment’ in the FASTA header and this resulted in 155 cI-VFP sequences.

The 155 cI-VFP sequences were organized by their virus families: Arenaviridae, Filoviridae, Orthomyxoviridae, Paramyxoviridae, Pneumoviridae, Coronaviridae and Retroviridae, according to the ICTV (International Committee on Taxonomy of Viruses) classification system (https://ictv.global/taxonomy/). Each Family was further divided using Genus information. Within a Genus, sequences were aligned to calculate sequence identity. If sequences were more than 75% identical, then only one of the sequences was chosen at random and retained in the dataset, thereby removing the over-represented viruses. This resulted in a final list of 65 sequences of cI-VFPs.

In this dataset (Supplementary table S1), 11 genera were represented by more than one sequence because these sequences were less than 75% identical: for instance, 9 Alphainfluenzavirus HA sequences (each belonging to a different class: H1, H2, H3, H4, H5, H7, H10, H12 and H13) were present. Overall, this dataset of 65 cI-VFPs comprises highly diverse proteins that are less than 25% identical to each other across the virus Families (Figure S1). Intra-Family average sequence identity ranges from 48% in Arenaviridae to 16% across Retroviridae.

#### Human fusion spTMPs

To assemble this dataset, we first searched by using the keyword “fusion” in Uniprot. The results were filtered by Reviewed sequences (3990 sequences), and then restricted to human sequences (model_organism=human) with “protein level” existence (1011 sequences). This sequence dataset included membrane fusion proteins, dynamin-like GTPases, nucleotide exchange factors, transcription factors, all of which directly or indirectly influence fusion in/of human cells. To isolate only human fusion proteins with a spTMH, the Membranome database (*45, 46*), a database of spTMPs from diverse model organisms, was queried (29^th^ July, 2024) using the 1011 sequences. This resulted in a list of 186 putative human fusion spTMP sequences. However, in addition to fusion proteins, this list contained receptors such as the bone morphogenetic protein (BMP) receptor that is useful in the process of bone fusion, cellular adhesion proteins that facilitate fusion, and receptors for viral fusion proteins such as TMPRSS2, dipeptidyl peptidase, etc. We tried to isolate the fusion proteins from this list by filtering using keywords such as “receptor”, or an enzyme classification code, but could not get a clean dataset. So, we turned to manual curation to filter the 186 spTMP sequences.

Two kinds of human fusion proteins have been well characterized, (exocytic/neuronal) vesicle fusion proteins, and cell-cell fusing syncytins (*5, 9*). The final human fusion protein dataset had 18 proteins: 16 SNARE proteins (*21*) that belong to the core vesicle fusion machinery (*47*), and both the human syncytin proteins, Syncytin 1 and 2 (*48*).

#### Human non-fusion spTMPs

A set of 65 human non-fusion spTMP sequences was curated from the Membranome spTMP database (*45, 46*) to compare with the cI-VFP dataset. These sequences were chosen manually to include functionally diverse proteins from all major biomembranes, specifically, plasma membrane, endoplasmic reticulum membrane, Golgi apparatus membrane, nuclear membrane, mitochondrial membranes and other endomembranes (e.g. lysosomes). It was important to ensure representation from different cellular membranes because these membranes differ in their thickness (*49*), and hydrophobic matching to membrane thickness can constrain the length of the TMHs of proteins native to that membrane (*35*). Additionally, the composition of this dataset (Supplementary Table S2) was chosen to match the global human spTMP membrane localization prediction (*50*).

We also tried to match the composition of the dataset to the functional classification of human spTMPs (*51*). Some of the proteins that we included are well-studied receptor tyrosine kinases (epidermal growth factor receptor (EGFR), insulin receptor (InsR)), endomembrane signaling proteins (stromal interaction protein 1 (STIM1)), cytoskeleton scaffolding/tethering proteins (emerin, nesprin-1, junctophilin), regulatory proteins of vesicle fusion (synaptotagmin), apoptosis regulators (BAX and Bcl2), etc. Some proteins in this list have TM domains that perform a passive structural/anchoring function (e.g. furin, glycophorin A, β-1,3-glucosyltransferase), whereas many others exist in a monomer-dimer equilibrium (e.g. Integrins) or self-assemble as multimers (e.g. phospholamban). Proteins with some experimental information on the TMH were included, when possible, in order to facilitate the interpretation of the TM length predictions. The final dataset of 65 proteins was highly diverse such that some sequences failed to align and the identity between aligned full-length proteins was only 12% on average.

### Transmembrane helix prediction

We used 11 well-known transmembrane prediction algorithms for determining the membrane spanning segment within each protein sequence (Figure 2A). All the algorithms were used at their default settings unless noted otherwise. The Kyte-Doolittle algorithm (*52*), which performs a hydropathy scan on a sliding window, was run from the ProtScale module on the Expasy server (https://web.expasy.org/protscale/) (*53*) using the default scanning window of 9 residues and an exponential weight variation model. Zhao and London’s TM tendency analysis (*54*), which includes a refinement to the hydrophobicity scale by incorporating information from known protein structures, was also performed on the Expasy server (https://web.expasy.org/protscale/) with settings identical to the Kyte-Doolittle model. HMMTOP (https://hmmtop.pbrg.hu/) (*55, 56*), Phobius (https://phobius.sbc.su.se/) (*58, 59*), and TMHMM (https://services.healthtech.dtu.dk/services/TMHMM-2.0/) (*57*), all of which use an implementation of hidden markov modelling to define a TM domain were used at their respective webservers. TOPCONS (*60*), which identifies a consensus from topology prediction algorithms, was used from its webserver (https://topcons.cbr.su.se/) (*61*). A local installation of MPEx (downloaded from https://blanco.biomol.uci.edu/mpex/) (*62*) was used in the hydropathy analysis mode, which creates a hydropathy plot based on an augmented White-Wimley scale (*63, 64*). This result is labelled MPEx in the text. Additionally, the implementation of ΔGPred on MPEx was used. ΔGPred uses experimentally-derived sequence requirements for TMH recognition by the Sec61 translocon (*65*) to predict TMHs. MEMSAT-SVM (*66*), a support vector machine based TM predictor was used at the PSIPRED workbench (https://bioinf.cs.ucl.ac.uk/psipred/) (*67*). SPLIT4 (*68*), which predicts TM domains by identifying basic charge clusters around hydropathic stretches of sequence was also run from its webserver (http://splitbioinf.pmfst.hr/split/4/). TMPred was run from a locally compiled version downloaded from https://github.com/rcanovas/TMPred. The ‘strongly preferred’ model was chosen as the result from TMPred. Literature was consulted to resolve the cases where more than one TM segment was predicted. The absolute C-terminal predicted segment was the correct prediction in all cI-VFPs other than a few Orthoretroviridae sequences (e.g. Walleye Dermal Sarcoma Virus), in which a part of the C-terminus (including the segment predicted as a TMH) is cleaved during the maturation phase of the fusion protein (*69*).

**Figure 2.**
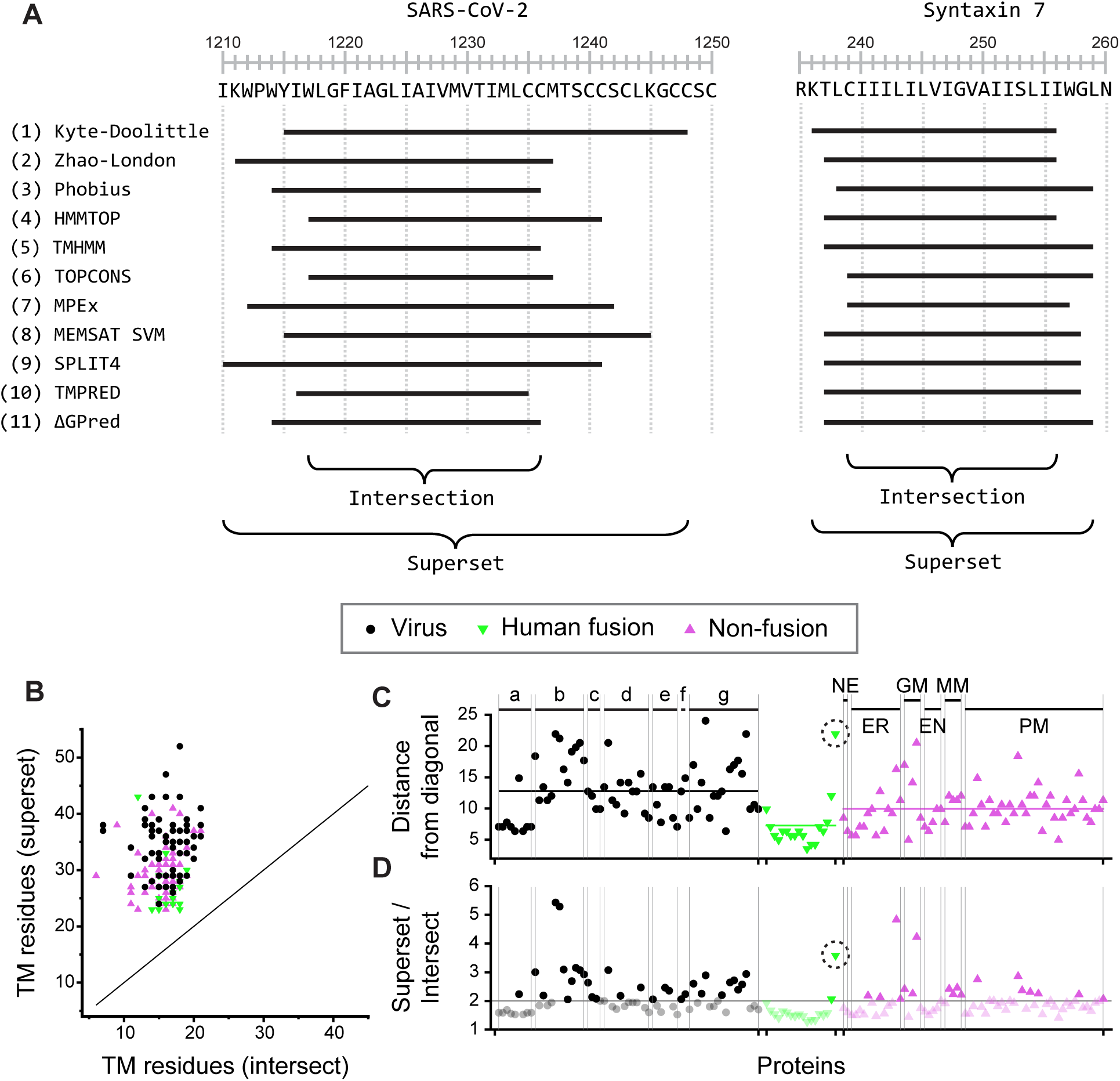
TM length prediction. (**A**) The 11 TMH prediction algorithms are listed on the left. Protein sequence and numbering follow Uniprot. Horizontal black lines adjacent to the prediction algorithm mark the predicted TMH sequence for that algorithm. The predicted TMH sequence common to all the algorithms: their intersection, and the sequence which is the superset of all the sequences are marked below the predictions. Left panel: SARS-CoV-2 spike protein. Right panel: Syntaxin 7, a vesicle fusion protein. When the TMH predictions across algorithms are similar (e.g. Syntaxin 7) then the Intersection length is more similar to the Superset length. (**B**) The superset length plotted versus the intersection length (both in number of residues) for the spTMHs: class I viral fusion proteins (black), the human fusion proteins (green) and human non-fusion proteins (pink). The diagonal is marked to show that, as expected, the superset length is always greater than the intersection length. (**C** and **D**) Color scheme as in (B). a-g shown above vertical divisions are virus families labelled as in Table S1. For the non-fusion proteins, the inhabited membranes are marked: Nucleus envelope: NE, endoplasmic reticulum membrane: ER, Golgi apparatus membrane: GM, other endomembranes: EN, mitochondrial membrane: MM, and plasma membrane: PM. (**C**) The perpendicular distance of each point in (B) from the diagonal. Colored horizontal lines mark the average values for the individual datasets and indicate that the viral TM-helices have higher variability across predictions. (**D**) Superset to intersection ratio for every protein in the dataset. A horizontal line is marked at 2 which is close to the ratio of the average value of the superset across the entire dataset (=31.3 residues) to the average value of the intersection (=15.9 residues). Syncytin 1, the human fusion protein outlier, is circled in both (C) and (D) and has spTMHs with properties similar to those of the virus fusion proteins.

Each algorithm predicted a putative TMH for all proteins in the dataset and so, most of the 148 spTMPs had 11 predicted spTMH sequences. Rarely, one or more of the programs would fail on a given spTMP sequence (Supplementary Table S3). Across the various algorithms, HMMTOP failed to predict a TMH 18 times out of 148, followed by TMHMM (9 times). SPLIT4 and Phobius both failed on two sequences each, and TOPCONS failed on one sequence. Rest of the algorithms worked for all the 148 spTMP sequences. The Arenaviridae Family had the most proteins for which more than one algorithm failed to successfully predict a TMH. In the human non-fusion dataset, spTMPs whose spTMHs were adjacent to the N-terminus of the protein caused the most failures. No prediction failed from the human fusion dataset.

### Analysis of TM lengths

Two parameters, superset and intersection (intersect), were calculated using the results of the TMH predictions. The superset of the TM predictions for a given spTMP is defined as the shortest continuous segment which contained all residues predicted as an spTMH by any of the 11 prediction algorithms (Figure 2A). The intersection is the consensus (or the longest) segment predicted by every prediction algorithm (Figure 2A). Distance from the diagonal in Figure 2B was calculated for each datapoint using the formula 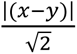 and is plotted in Figure 2C. Mean length for each spTMH (Figures 3 and S2) was calculated by averaging the lengths (number of residues) of the TMH obtained from the 11 prediction algorithms. The standard deviation, 𝜎, is calculated using 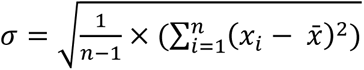, where *x_i_* is an individual point (e.g. mean spTMH length of a given spTMP in Figure 3 and S2) in the dataset, 𝑥̄ is the average of 𝑥_𝑖_, and 𝑛 is the number of points in the dataset. The standard error (Figure 3 insets) is 𝜎/√𝑛. Significance was calculated by the Student’s t-test with a Welch correction (equal variance not assumed).

**Figure 3.**
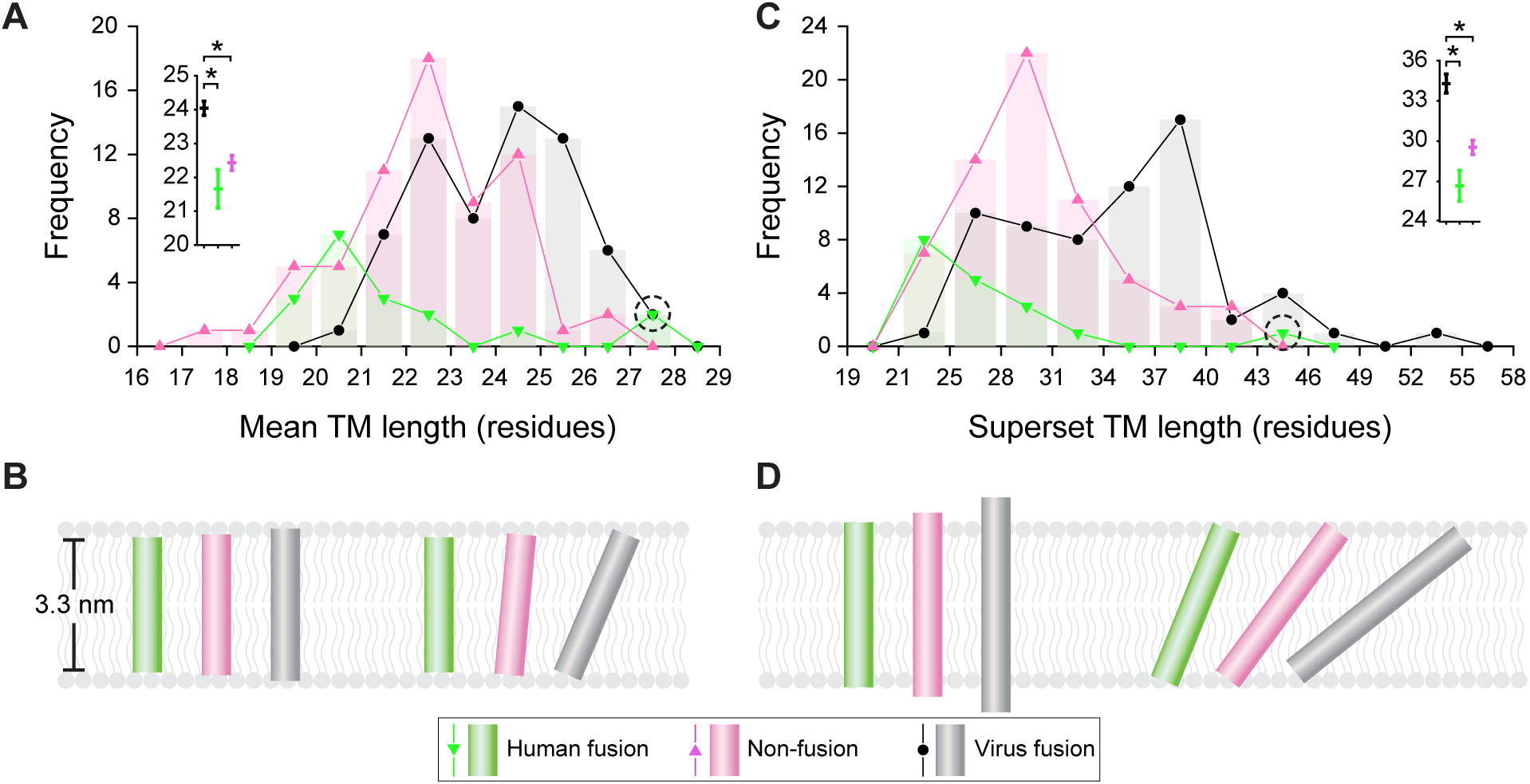
Analysis of predicted TMH lengths. (**A**, **C**) Histograms of (A) the mean predicted TM length (number of residues) and (C) the superset TM length (number of residues) of the three sets of proteins: cI-VFP (black), human fusion (green), and human non-fusion (pink). Each bar shows the number of proteins that have a given length. The centers of the bars are marked and joined by a line to guide the eye. The insets show the average of the lengths ± the standard error for each dataset in the same color as the data. *indicates significance at p<0.001. Syncytin 1, the human fusion protein outlier, is circled. (**B**, **D**) The heights of the shaded rectangles depict the heights of helices. ((B): human fusion = 3.2 nm, non-fusion = 3.4 nm, cI-VFP = 3.6 nm; (D): human fusion = 4.1 nm, non-fusion = 4.4 nm, cI-VFP = 5.2 nm). These were calculated in (B) using the average of the mean TM length in number of residues (see inset of (A): human fusion = 21.7, non-fusion = 22.4, cI-VFP = 24) and in (D) using the average of the superset TM length in number of residues (inset of (C): human fusion = 26.7, non-fusion = 29.5, cI-VFP = 34.3), and multiplying each by 0.15 nm (ideal rise per residue of an α-helix). Rectangles are colored as in (A, C). A typical biomembrane of 3.3 nm hydrophobic thickness is shown in pale gray. The helices are placed perpendicular to the membrane on the left in (B, D) and it can be seen that the longer helices protrude outside the membrane. The same helices are tilted to fit the membrane on the right. (B) The average (of the mean) viral TMH has a tilt of ∼23° with respect to the membrane normal. The non-fusion helix has a tilt of ∼5°. (D) The longer average superset helices are more tilted. The 34.3 residue long viral TMH has a tilt of 52°.

### Analysis of amino acid distributions

The composition of the sequences (Figure S6) was calculated using the equation 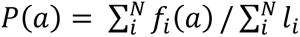 where, 𝑃(𝑎) is the probability of occurrence of any amino acid 𝑎 in all the sequences in the dataset, 𝑓_𝑖_ (𝑎) is the frequency of occurrence of an amino acid in the 𝑖^𝑡ℎ^sequence, 𝑁 is the number of sequences in the dataset, and 𝑙_𝑖_ is the length of the 𝑖^𝑡ℎ^ sequence. For example, the total number of Ile residues in the complete 148 spTMP dataset=5839 and the total residues in the dataset=103807. Therefore 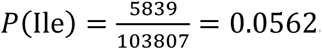.The percentage occurrence for an amino acid, 𝑎, is 𝑃(𝑎) × 100. To calculate the percentage occurrence of the TM region (Figure S6B), we used the superset segment of each spTMH plus five residues (pad) on either side to calculate 𝑓_𝑖_ (𝑎). To calculate propensity, the percentage occurrence in the superset+pads was divided by percent occurrence in the full-length 148 spTMP dataset.

## RESULTS

### Prediction of single pass transmembrane helix (spTMH) segments

Due to limited experimental spTMH data, we used computational predictors to infer the TMH segments of the proteins in the three curated spTMH datasets: class I viral fusion proteins (cI-VFPs), human fusion single pass transmembrane proteins (spTMPs), and human non-fusion spTMPs. There are multiple well-known algorithms/predictors based on different properties/methodologies that can predict spTMH segments, and we analyzed the spTMH predictions from eleven such algorithms (Methods). We calculated the mean predicted spTMH length (number of residues) for a given spTMP by averaging over the length predictions from all the algorithms. The combined prediction from within each dataset (Figure S2) shows that the average of the mean spTMH length (*l* in number of residues) for the spTMH predictions of the cI-VFPs (𝑙 = 24, σ = 1.7) > human non-fusion spTMPs (𝑙 = 22.4, σ = 1.8) > human fusion spTMPs (𝑙 = 21.7, σ = 2.3), where σ is the standard deviation of the mean spTMH lengths in a given dataset.

### Variability in spTMH predictions

We had previously observed that different algorithms predicted different spTMH segments for the SARS-CoV-2 spike protein (Figure 2A) (*19*). Examining the predictions for many spTMHs from the different algorithms (Figures S3 and S4; also see Methods) we find the following: Some predictors have a low standard deviation across the combined data from the three datasets (Figure S3) because their prediction lengths are additionally constrained. For example, TOPCONS: sliding window of 21 residues used in the webserver (*61*); ΔGpred: training on the translocation of model peptides across the Sec61 translocon (*65*); TMHMM: known to produce results with a high representation of 20 and 23 residue TMHs (*59*). Most other algorithms exhibit a higher prediction standard deviation (Figure S3) and produce spTMH length distributions centered at ∼20-25 residues (Figure S4). Generally, the hydropathy prediction algorithms (Kyte-Doolittle, Zhao-London, MPEx) have the most variable spTMH length predictions (Figure S3 and S4).

In summary, for any given spTMP, there is prediction variability, i.e., different TM prediction algorithms give different results for the spTMH segments and lengths. Thus, the use of multiple predictors followed by an examination of the predictions may be needed before deciding on an spTMH segment. We next compare the spTMH prediction variability between the three curated datasets: cI-VFP, human fusion spTMPs and human non-fusion spTMPs.

### cI-VFP spTMHs have a high prediction variability

In order to understand the variability in spTMH predictions for a given protein across the eleven algorithms, we constructed two parameters (Figure 2A): the superset length which is the minimum continuous segment containing all the residues from every prediction, and the intersection length (intersect) which is the maximal segment containing residues present in all the predictions. By definition, the superset length ≥ intersect length, with the two being equal only if the predictions across algorithms are identical. So, all the proteins lie on the left side of the diagonal in the superset vs intersect plot (Figure 2B). The larger the variability in the predictions, the larger the difference between the superset length and the intersect length. In Figure 2B, the human fusion spTMPs cluster close to the diagonal, the non-fusion spTMPs and the cI-VFPs being further away from it. The distance of each spTMP from the diagonal confirms this trend (Figure 2C). This distance is proportional to the length difference between the superset and the intersect (see Methods), and is thus a direct readout of variability. As expected, the human fusion spTMPs are the closest to the diagonal, being on average 7.3 residues away, with average of superset length−intersection length (AV)=10.3 residues, while the non-fusion proteins are 9.9 residues away (AV=14 residues) and the cI-VFPs, are the furthest from the diagonal being on an average 12.8 residues away (AV=18.1 residues) (Figure 2C). So, the cI-VFPs have the largest variability in spTMH segment predictions.

We also calculated the ratio of superset to intersect of each spTMH and compared it to the ratio of average superset length across the three datasets combined (=31.3 residues) to the average intersect length (=15.9) which is ≈2 (Figure 2D). The spTMHs of most cI-VFPs (one of the exceptions being Arenaviridae discussed later) have a high superset to intersect ratio. Two bat CoVs HKU4 and HKU5 have unusually high superset to intersect ratios because of short intersect (7 residues) and long superset (38 and 37, respectively) lengths. This occurs because a single algorithm (TOPCONS) predicts an spTMH segment which has a low overlap with the consensus (intersect) segment of the other algorithms. Overall, 48% of cI-VFPs had a superset to intersect ratio > 2, while only 28% of human non-fusion spTMPs have a ratio > 2. Only two human fusion spTMPs, the retrovirus-derived glycoproteins, syncytins have a ratio >2 (Figure 2D). One of these, Syncytin 1 (superset length=43, intersect=12), stands out from the other human fusion spTMPs and has a distance from the diagonal (Figure 2B) of 22 residues (Figure 2C) and a superset to intersect ratio of 3.6 (Figure 2D). The syncytins, their origin and function will be discussed in a subsequent section.

The more broadly studied human non-fusion spTMPs have superset to intersect ratios ranging from those similar to human fusion spTMPs (low) to those similar to cI-VFPs (high). Given this, we examine the properties of non-fusion spTMPs with high and low superset to intersect ratios separately, and infer the potential properties of the two sets of fusion spTMPs by comparison. Specifically, spTMPs with dynamic spTMHs (*30, 70*) (e.g. Cadherin 1, Integrin α1, Integrin β3, Toll-like receptor 1 (TLR1), T cell receptor component CD4, Erlin-1, Chondrolectin, and CD8) have a superset to intersect ratio of ≥ 2. In contrast, spTMPs that undergo homodimerization (e.g. Epidermal Growth Factor Receptor (EGFR), Insulin receptor (InsR), Vascular Endothelial Growth Factor Receptor 2 (VEGFR 2), and Erythropoietin receptor (EpoR)) have a superset to intersect ratio < 2. We note in passing that the predictions were inconsistent across algorithms for some non-fusion spTMPs from the Golgi and ER membranes with some algorithms predicting the spTMH-adjacent signal peptide as the spTMH. This caused an unusually high superset to intersect ratio.

Overall, the cI-VFPs have the highest variability in spTMH prediction. The non-fusion spTMHs with high variability are known to be dynamic and so, the high cI-VFP prediction variability may also imply dynamic spTMHs. We next examine the average length of the predicted spTMH segments.

### cI-VFPs have longer TM helices

We calculated the mean predicted spTMH length for a given spTMP by averaging over the length predictions (number of residues in the predicted spTMH for a given algorithm) from all the algorithms. Histograms of these mean lengths are plotted for each of the three datasets (Figures 3A and S2) and these show that the cI-VFPs have the longest spTMHs, followed by the human non-fusion spTMPs, and the spTMHs of the human fusion spTMPs are the shortest (Figure 3B). Thus, the variability in spTMH prediction correlates with the mean length of the spTMHs.

As stated in the previous sections, there are some spTMHs with unusually large superset lengths because different algorithms predict overlapping but distinct protein segments as the spTMH. Given this, we also plotted the histograms of the superset lengths of the three datasets (Figure 3C), and find that the superset spTMH distributions follow the same trends as the mean spTMH distribution. Overall, with the exception of the Arenaviridae and the Ebolavirus fusion proteins (discussed later), whose mean predicted spTMH lengths of 22.1 residues are similar to the average length of non-fusion spTMHs (22.4 residues), the mean predicted TM lengths of virus Families are longer (Figure S2).

The average of the mean spTMH lengths for human fusion spTMPs was 21.7 residues (Figure S2), which is just long enough to traverse the hydrophobic interior of a lipid bilayer that is ∼3.3 nm thick (thickness of a model lipid bilayer (*71–73*), assuming an ideal α-helix with 3.6 residues per turn, and a canonical pitch of 0.54 nm (*34, 74, 75*)). Since most vesicle fusion proteins inhabit vesicles derived from the ER or the Golgi membrane, which are among the thinner biological membranes, the spTMHs of such vesicle fusion proteins are, as expected, shorter and almost perpendicular to the membrane (Figure 3B). Both the syncytins (discussed later) perform cell-cell fusion, and have a mean spTMH length of 27 residues which is closer to that of the cI-VFP spTMHs. Thus, their spTMH is longer than the average spTMH of human vesicle fusion proteins by ∼5.3 residues, or roughly 1.5 helical turns. This would require them to tilt even in the thicker plasma membrane (Figure 3D).

Amongst non-fusion spTMPs, nuclear envelope residing Nesprin-1 and Emerin have a mean spTMH length of 22.1 and 20.3 residues, respectively. ER membrane resident spTMPs like STIM1, phospholamban, and junctophilin-1, have a mean spTMH lengths of 21.3 residues. Golgi and endosomal membrane localized proteins: Golgi membrane protein 1, Synaptotagmin-1, Lysosome-associated membrane glycoprotein 1 (LAMP1), LAMP3 all have mean spTMH lengths of 22.6 residues. Plasma membrane localized Glycophorin A (GpA), VEGFR 2, InsR, Prolactin receptor (PrlR), Platelet-Derived Growth Factor Receptor β (PDGFR β), EGFR, EpoR, Integrin β3, Angiotensin-converting enzyme 2 (ACE-2), have mean spTMH lengths of 23.2 residues. Further, known tilted TMHs (Integrin β3 and α1) are slightly longer with a mean spTMH length of 24.3 residues, while anchoring TMHs (e.g. ACE-2, average length 22.4 residues; GpA, 22.3 residues) are shorter. So, the mean spTMH length for non-fusion proteins seems organelle-specific and correlated with the thickness of the native membrane for the spTMP (Figure S2). Additionally, the mean lengths of known tilted spTMHs were slightly longer, and the variability in the prediction of the spTMH (reflected in, for instance, the superset to intersect length ratio) seems to correlate with the type of dynamics in the membrane as discussed in the previous section.

spTMH prediction variability need not be correlated with predicted spTMH length. However, we find that both prediction variability as well as average spTMH length are lowest for the human fusion spTMPs, followed by the human non-fusion spTMPs. Both, the prediction variability and the average spTMH length are the highest for cI-VFPs.

## DISCUSSION

### Possible reasons for the variability in the cI-VFP spTMH predictions

The cI-VFP spTMH predictions had the highest variability across the eleven TMH prediction algorithms (Figure 2). A possible reason is the residue signals that determine the spTMH segment are less apparent in the cI-VFPs. The different cI-VFP spTMH sequences were not easy to align (Figure S5), and so it is likely that there isn’t one single amino acid or factor responsible for the uncertainty in the spTMH prediction for all cI-VFPs. In order to test this, we analyzed the amino acid composition of the cI-VFPs.

The overall compositions of the entire Uniprot database (*43*), the Membranome spTMP database (*45, 46*), and the combined 148 sequence dataset used here were similar (Figure S6A). This implies that the membrane-spanning regions of the databases are small and the major contributor to the amino acid composition is the soluble region of the proteins. As expected, the amino acid composition of the TM region of the combined 148 sequence dataset is distinct from that of the full length spTMPs (Figure S6B). Additionally, there are differences in the propensities of two amino acid residues between the TM regions of the three datasets (Figure S6C). Specifically, there is a higher propensity of Cys residues in the cI-VFP TM region, and there is a higher propensity of Trp in both the fusion protein datasets compared to the non-fusion dataset. We next examine residue propensities in specific virus families.

In coronavirus (CoV) cI-VFPs, there are several aromatic residues on the N-terminal side of the spTMH (*76, 77*), a region we recently termed TRACS (Tryptophan Rich Aromatic Conserved Stretch) due to an enrichment of conserved Trp residues (*19, 78*). These conserved Trp residues are important for membrane fusion (*79–81*). Additionally, there is a Cys-rich weakly polar stretch, also comprising Met and Ser/Thr residues on the C-terminal side of the spTMH. Although the functional role of the other residues is unexplored, palmitoylation at selected Cys residues improves membrane fusion (*82*). The spTMH predictions for a given CoV may not agree because of differing weights given to the terminating residues (the long aromatic stretch at the N-terminus and the weakly polar residues at the C-terminus of the spTMH) in different algorithms.

Some retroviruses are known to have a basic Arg residue near the center of the spTMH (*83, 84*). This Arg is accommodated in the hydrophobic membrane interior through hydrogen bonding with adjacent backbone residues and a water molecule (*85*). This unusual Arg residue could confound both hydropathy-based predictions, as well as predictions based on non-retroviral training data.

A long hydrophobic TM sequence should, on an average, result in a longer spTMH prediction. The presence of qualitatively different length constraints (such as a sliding window in the prediction algorithm) may result in different segments of the hydrophobic stretch being predicted as the spTMH by different algorithms, leading to a positive correlation between spTMH length and variability. In practice, cI-VFP prediction variability is likely due to a long hydrophobic segment as well as due to unclear termination signatures. We next examine the possible effects of the longer hydrophobic stretches of spTMHs on cI-VFP function.

### Hydrophobic-mismatch to the lipid membrane may induce spTMH dynamics

Viral envelopes originate from different host cell membranes, and are thus, presumably, no thicker than the plasma membrane (also see Table S2). The long average (of the mean) spTMHs (∼24 residues) and an even longer superset spTMH segment (∼35 residues) in cI-VFPs implies that these spTMHs will be longer than the nominal length needed to traverse the viral envelope in which they reside (Figure 3B and 3D).

The behavior of spTMHs is particularly sensitive to changes in length. For example, a single extra residue can exert differential self-association in SNARE proteins (*86*). So, the length of the cI-VFP spTMHs should affect their assembly and dynamics, especially since they are on an average (length of the mean), two residues longer than non-fusion protein spTMHs, whose TMH lengths correlate with the thickness of the membrane that they are located in (*35, 87*). In fact, the hydrophobic-mismatched cI-VFP spTMHs would exhibit conformational dynamics to minimize the exposure of their hydrophobic surface as is seen in other membrane proteins (*33, 36, 88*) and under model conditions (*37*). Experiments (*89*) and molecular dynamics (MD) simulations show that this hydrophobic mismatch manifests itself by tilting, bobbing and/or bending of the TMH in HIV-1 (*39*) and SARS-CoV-2 (*19*). Also, the entire length of the spTMH may not remain helical at all times, becoming partially helical in the trimeric pre-fusion state (Figure 1A), and more helical and tilted, when separated (*19*). Such spTMH dynamics could lead to the disassembly of the TMD (composed of 3 spTMHs; Figure 1), and destabilization of the cI-VFP trimer, thereby, improving the kinetics of conversion to the post-fusion state.

cI-VFP spTMH conformational dynamics can also induce local deformations in the envelope (*19, 39*). If the average predicted superset cI-VFP spTMH segment tilts (Figure 3B and 3D), it can destabilize an area of about 6.2 nm^2^ at the surface of the envelope (a 3.6 nm cI-VFP spTMH with a tilt of 23° from the membrane normal (Figure 3B) forms a cone of radius 3.6 × sin 23 ≈ 1.4 nm and a base area of 𝜋𝜋 × (1.4)^2^ ≈ 6.2 nm^2^). Such perturbations may promote fusion of the destabilized envelope with a proximal host bilayer.

### Arenaviridae and Filoviridae: cI-VFPs with shorter spTMHs

In this section, we discuss two virus families with shorter than expected spTMHs, and potential functional reasons for these. The predictions showed Filoviridae (same family as Ebolavirus) cI-VFPs to have their TMHs at the absolute C-terminus of the protein. So, similar to tail-anchored membrane proteins such as SNAREs, they have an almost non-existent cytoplasmic tail (*86*). Thus, there is little end prediction variability at the C-terminus of the spTMH, so, the mean of the predicted spTMHs is shorter. There is also some evidence that the Filoviridae cI-VFP spTMHs do not behave in a manner similar to those of other class I enveloped viruses. For example, unlike in SARS-CoV-2 and HIV-1, NMR spectroscopy in diverse lipid environments reported Ebola virus cI-VFP spTMHs to be monomeric (*90*). Additionally, the role of the Ebola virus spTMH in viral fusion is still unclear, because while mutating two Gly residues in the spTMH has little effect on viral entry (*91*), mutating the same residues may facilitate viral fusion via a cholesterol dependent mechanism (*92*).

Unlike the trimeric spTMH organization of the TMD in other cI-VFPs (Figure 1A), the Arenaviridae family members, Lassa virus (*93*) and Lujo virus (*94*) have a 6 helix TMD in the pre-fusion state. This includes not only the three C-terminal spTMHs, but also three N-terminal (of the full length cI-VFP) stable signal peptide (SSP) helices. The SSP-TMD interaction is important for subsequent pH-change triggered Arenaviridae membrane fusion (*95*). Also, it was recently shown that the SSP undergoes conformational dynamics in response to a pH change (*96*). So, it is possible that the SSP helix takes over the dynamic role performed by the spTMHs from other cI-VFP families. Incidentally, the Lassa virus cI-VFP has also been suggested to possess viroporin activity (*97*), a function that is not usually associated with fusion proteins. Overall, the spTMHs of the Ebolaviridae and Arenaviridae cI-VFPs may be shorter because the cI-VFPs have non-canonical structures and perform non-canonical functions.

### A comparison between cI-VFP and human fusion spTMHs

It was believed that the spTMHs of vesicle fusion spTMPs play an anchoring role, and do not perform large conformational dynamics (*98*). However, replacing the spTMH in SNARE proteins with lipid anchors has been shown to both support fusion (*98*), or have no effect on it (*99*). Unlike the “one protein does it all” cI-VFPs, SNAREs are part of protein complexes, and the contradictory effects of their spTMHs (*100*) could be due to the contribution of proteins other than SNAREs (*5*).

Contrasting evidence notwithstanding, there seems to be an emerging role for conformational dynamics in the vesicle fusion proteins’ spTMHs. For instance, data from both full-length vesicle fusion spTMPs (*101*) and isolated TMHs (*102*) shows that TMH flexibility is important for spTMP ability to fuse vesicular membranes. For instance, residues that decrease helicity such as Pro, Gly, and even Val, are positively correlated with fusability, whereas a poly-Leu spTMH impaired membrane fusion (*102, 103*). Vesicle fusion spTMHs may, in some cases, be even shorter than the membrane, inducing membrane dimples (*86*), and lipids that promote membrane curvature may complement spTMH conformational dynamics (*101*). Finally, monomer-dimer equilibria of some spTMHs can modulate vesicle fusion (*104*). Thus, vesicle fusion protein spTMH dynamics may also promote fusion, but these dynamics may be more helicity-dependent and local, compared to the larger scale bobbing, tilting and bending movements that are possible due to the longer cI-VFP spTMHs.

### Syncytins: human fusion spTMPs with longer spTMHs

The Syncytin 1 sequence from the human fusion protein spTMH dataset has both prediction variability (distance from the diagonal of the superset-intersect plot ∼22 residues: Figure 2C, and superset to intersect ratio of ∼3.6: Figure 2D), and average TM length (27 residues) similar to that of cI-VFP spTMHs. This is not surprising because the two human syncytins (Syncytin 1 and Syncytin 2) are encoded by endogenous retroviral envelope glycoprotein (captured cI-VFP from an ancient retroviral infection) genes (*105–107*). Syncytins are known to fuse the cells that form the placenta, and have been instrumental in the diversification of mammals from their egg laying ancestors (*48*). Although Syncytin 2 has a lower spTMH prediction variability (superset to intersect ratio just above 2) than Syncytin 1, their mean spTMH lengths are similar. Using a phylogenetic comparison of endogenous retroviruses with their extant relatives (*108*), we find that the average of mean spTMH lengths of 8 Syncytin proteins from various mammals (25.5 residues; Supplementary information) is similar to those of cI-VFP spTMHs of closely related extant retroviruses, namely, moloney murine leukemia virus (MoMLV; mean spTMH = 25.8 residues), mouse mammary tumor virus (MMTV 23.6 residues) and mason-pfizer monkey virus (MPMV 26.1 residues). So, the spTMH length of endogenous retrovirus fusogenic proteins correlates better to sequences of their originating viruses than to their present host membranes. This data reinforces our hypothesis that the longer cI-VFP spTMHs are important for membrane fusion.

### Targeting the cI-VFP spTMHs to modulate viral fusion

The long cI-VFP spTMHs have a hydrophobic mismatch with biological membranes and this is likely to make the cI-VFPs dynamic. The presence of such dynamics could be tested experimentally using NMR (*109*) or MD simulations (*19, 39*). Such dynamics may enable viral fusion by improving the pre-to post-fusion conformational transition of the cI-VFPs (Figure 1) or by locally destabilizing the membrane. So, modulating the spTMH dynamics may reduce viral fusion, and in turn viral infection. For example, replacing the TMD by a trimerizing domain is a trusted strategy to stabilize cI-VFPs in their pre-fusion state (*110, 111*). Additionally, regions within the cI-VFP, which have sequence and structure conservation (*4, 6, 112*), such as some spTMHs, can be targets for broad viral Family therapeutic interventions. This strategy of targeting conserved regions has recently been explored to lock the S2 region of the SARS-CoV-2 spike into its pre-fusion state to make pan-Coronaviridae vaccines (*110, 111*).

Lipid partitioning peptides that stabilize one conformation over the others have become a reality for cytokine receptors that are anchored to the plasma membrane with an spTMH (*113*). So, designing peptidic inhibitors of viral fusion by targeting the understudied spTMH dynamics is possible. Modulating the spTMH dynamics may also help stabilize full length cI-VFP in the pre-fusion conformation so as to serve as an effective antigen for vaccine production (*16*). Given the structure of the TMD in the post-fusion structure of the spike protein of SARS-CoV-2 (*78, 114*), we suggest the addition of the fusion peptides (Figure 1) either to the membrane or to the C-terminus of the cI-VFPs to obtain more such structures.

## CONCLUSION

The transmembrane domain (TMD) of class I viral fusion proteins (cI-VFPs) is comprised of three single-pass transmembrane helices (spTMHs). Not much is known about the contributions of the cI-VFP TMD to viral fusion. By comparing the lengths of the spTMHs from carefully curated datasets of (1) cI-VFPs, and spTMH-containing (2) human fusion, and (3) human non-fusion proteins, we show that on average the cI-VFP spTMHs are longer than human fusion spTMHs, which are known to be a better fit to biological membranes. Since our cI-VFP dataset consists of proteins from diverse enveloped virus families, we conclude that the long spTMH lengths are preserved across the dataset because they enable function. Specifically, the longer cI-VFP spTMHs create a hydrophobic mismatch with the membrane. This mismatch could lead to spTMH dynamics, which would consequently facilitate the cI-VFP pre-fusion to post-fusion conformational transition. The mismatch could also lead to membrane dynamics and destabilization. Both these effects are likely to promote membrane fusion. There are two endogenous retroviral envelope (cI-VFPs of retroviruses) proteins in our human fusion protein dataset, the syncytins, which are remnants of ancient retroviruses that have become integrated into the human genome. In further support of the functional role of the long cI-VFP spTMHs, we find that despite being part of the human genome for millions of years, the syncytin spTMH lengths are similar to those of closely related cI-VFPs. We expect that an understanding of cI-VFP spTMH dynamics will aid the design of new anti-virals that target the fusion process.

## Acknowledgements

This work was funded by the Department of Atomic Energy, Government of India through the Tata Institute of Fundamental Research from Project Identification No. RTI 4006. SL was supported with a postdoctoral fellowship through the Simons Foundation (Grant No. 287975). SG was supported in part by the Simons Foundation (Grant No. 287975), a grant from the National Supercomputing Mission (NSM) through the grant MeitY/R\&D/HPC/2(1)/2014 and SERB Grant: CRG/2021/004754.

## Data availability

All the relevant datasets associated with this manuscript will be provided as Supporting Information files upon manuscript publication.

## Supplementary Information

The supporting information contains supplementary tables S1 to S3 and supplementary figures S1 to S6

