## Supplementary Information for "The longer transmembrane helices of class I viral fusion proteins may facilitate viral fusion"

### for

| | Virus Family | Number of proteins |
|---|---|---|
| a | Arenaviridae | 9 |
| b | Coronaviridae | 13 |
| c | Filoviridae | 4 |
| d | Orthomyxoviridae | 12 |
| e | Paramyxoviridae | 7 |
| f | Pneumoviridae | 2 |
| g | Retroviridae | 18 |

**Table S1.** Number of cI-VFPs in each virus family (in alphabetical order) in our dataset.

| Location | Number of proteins chosen | Percentage of proteins (%) | Percentage previously reported (%) |
|---|---|---|---|
| Plasma membrane (PM) | 35 | 53.9 | 57 |
| Endoplasmic reticulum membrane (ER) | 13 | 20 | 13 |
| Golgi apparatus membrane (GM) | 5 | 7.7 | 9 |
| Mitochondrial membrane (inner and outer) (MM) | 5 | 7.7 | 6 |
| Nuclear envelope (NE) | 2 | 3.0 | 2 |
| Other endomembranes (EN) | 5 | 7.7 | 5 |
| Undefined | - | - | 8 |

Table S2. Membrane localization of the human non-fusion spTMP dataset used here (localization data from Uniprot) compared to the human spTMP proteome (data from Pogozheva and Lomize, BBA-Biomembr, 2018). Three proteins in our human non-fusion dataset had an undefined localization. These were reassigned to membranes as follows: The subcellular location of two of the proteins classified as undefined in our dataset had a Uniprot annotation of "Membrane". These two (Uniprot IDs: Q6UW88 and P60606) were assigned to the PM, based on their Uniprot Function Annotation. The third protein, Uniprot ID: O94985, was assigned to the ER membrane based on its published function (Uniprot Annotation) as a molecular adapter in anterograde transport. Thus, the final dataset (first two columns) had no undefined/ unassigned proteins. Reported membrane thicknesses are PM: 4.25 ± 0.03 nm, GM: 3.95 ± 0.04 nm, ER: 3.75 ± 0.04 nm, (Mitra et al., 2004, PNAS). Interpreting the thickness of the membrane hydrophobic core from these measured lengths is difficult and so, we use only the trend of biological membrane thickness in this study.

| | class I viral fusion | human fusion | human non-fusion | total fails |
|---|---|---|---|---|
| TOPCONS | 0 | 0 | 1 | 1 |
| TMHMM | 6 | 0 | 3 | 9 |
| ΔGPred | 0 | 0 | 0 | 0 |
| Phobius | 1 | 0 | 1 | 2 |
| TMPRED | 0 | 0 | 0 | 0 |
| HMMTOP | 13 | 0 | 5 | 18 |
| SPLIT4 | 0 | 0 | 2 | 2 |
| MEMSAT-SVM | 0 | 0 | 0 | 0 |
| Zhao-London | 0 | 0 | 0 | 0 |
| Kyte-Doolittle | 0 | 0 | 0 | 0 |
| MPEx | 0 | 0 | 0 | 0 |

**Table S3.** Number of sequences from the three datasets that did not produce a result with a given prediction program/algorithm. The algorithms are ordered as in Fig. S4.

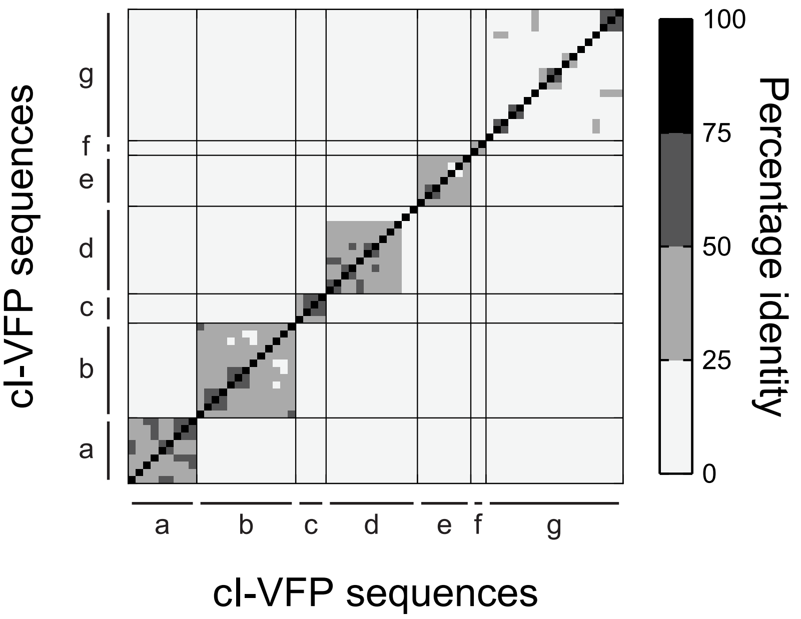


**Figure S1: Percentage identity matrix.** Pairwise percentage identity matrix of all the full-length class I viral fusion proteins (cI-VFPs). (a-g) the different virus families (as in Table S1). a: Arenaviridae, b: Coronaviridae, c: Filoviridae, d: Orthomyxoviridae, e: Paramyxoviridae, f: Pneumoviridae, g: Retroviridae.

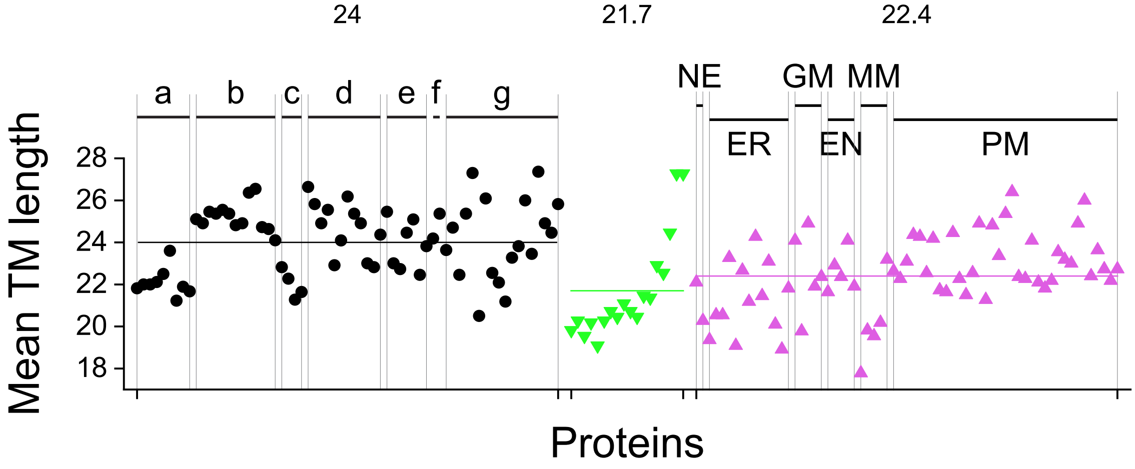


**Figure S2. Mean TMH length** (in number of residues) for every protein in the three datasets: cl-VFP (black), human fusion (green) and human non-fusion (pink). Colored horizontal lines mark the average values for the individual datasets with the value shown above the dataset. The viral TM-helices are on average longer than human fusion and non-fusion proteins. The two rightmost human fusion proteins are the syncytins, and they have lengths similar to those of cl-VFPs. Labels same as in Figure 2C. a-g shown above vertical divisions are virus families labelled as in Table S1. For the non-fusion proteins, the inhabited membranes are marked. Nucleus envelope: NE, endoplasmic reticulum membrane: ER, Golgi apparatus membrane: GM, other endomembranes: EN, mitochondrial membrane: MM, and plasma membrane: PM.

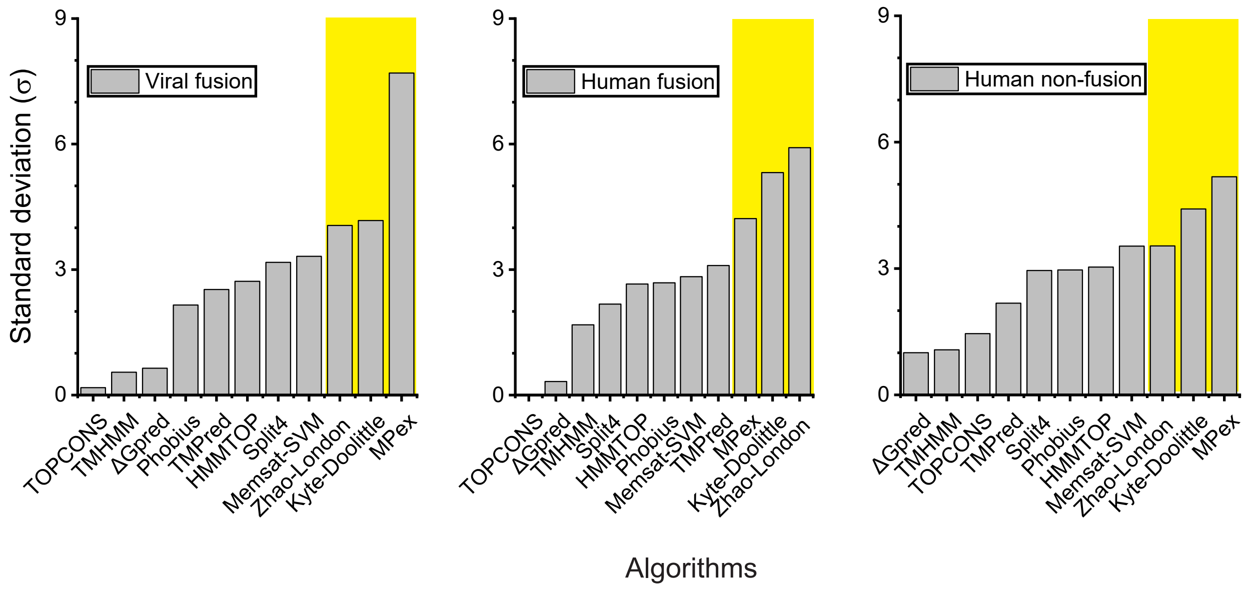


**Figure S3: Standard deviations in the predicted TMH length for each algorithm across the three datasets.** The algorithms are shown in ascending order of their standard deviation. The three algorithms with the highest standard deviation are highlighted in yellow, and are the same algorithms across the three datasets. Also, across datasets, the set of three algorithms that have the lowest standard deviation are the same.

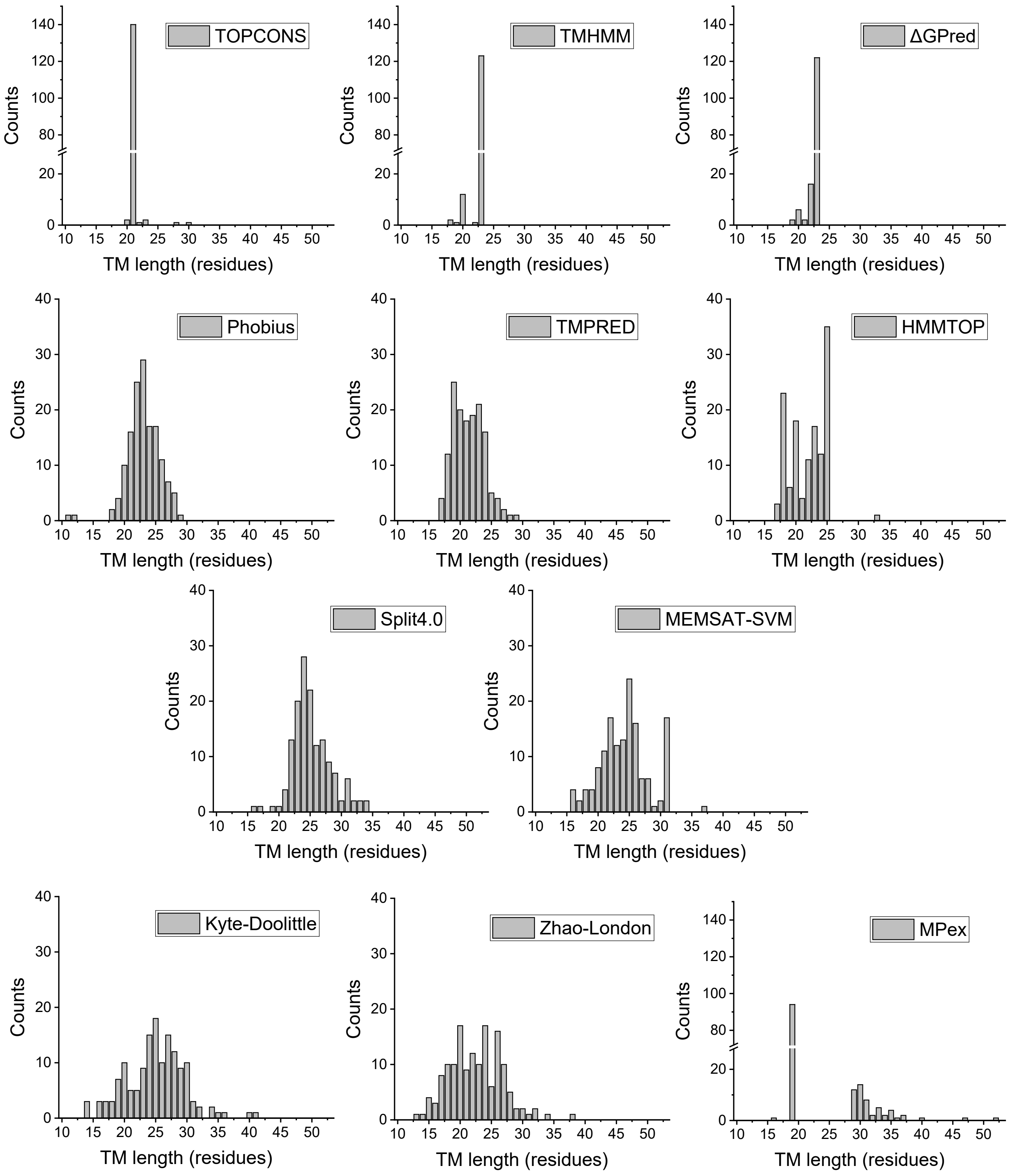


**Figure S4. Histograms of TMH prediction lengths for the entire 148 sequence dataset based on algorithms.** The algorithms are shown in ascending order of their standard deviation in viral fusion proteins (see Fig. S3, left panel).

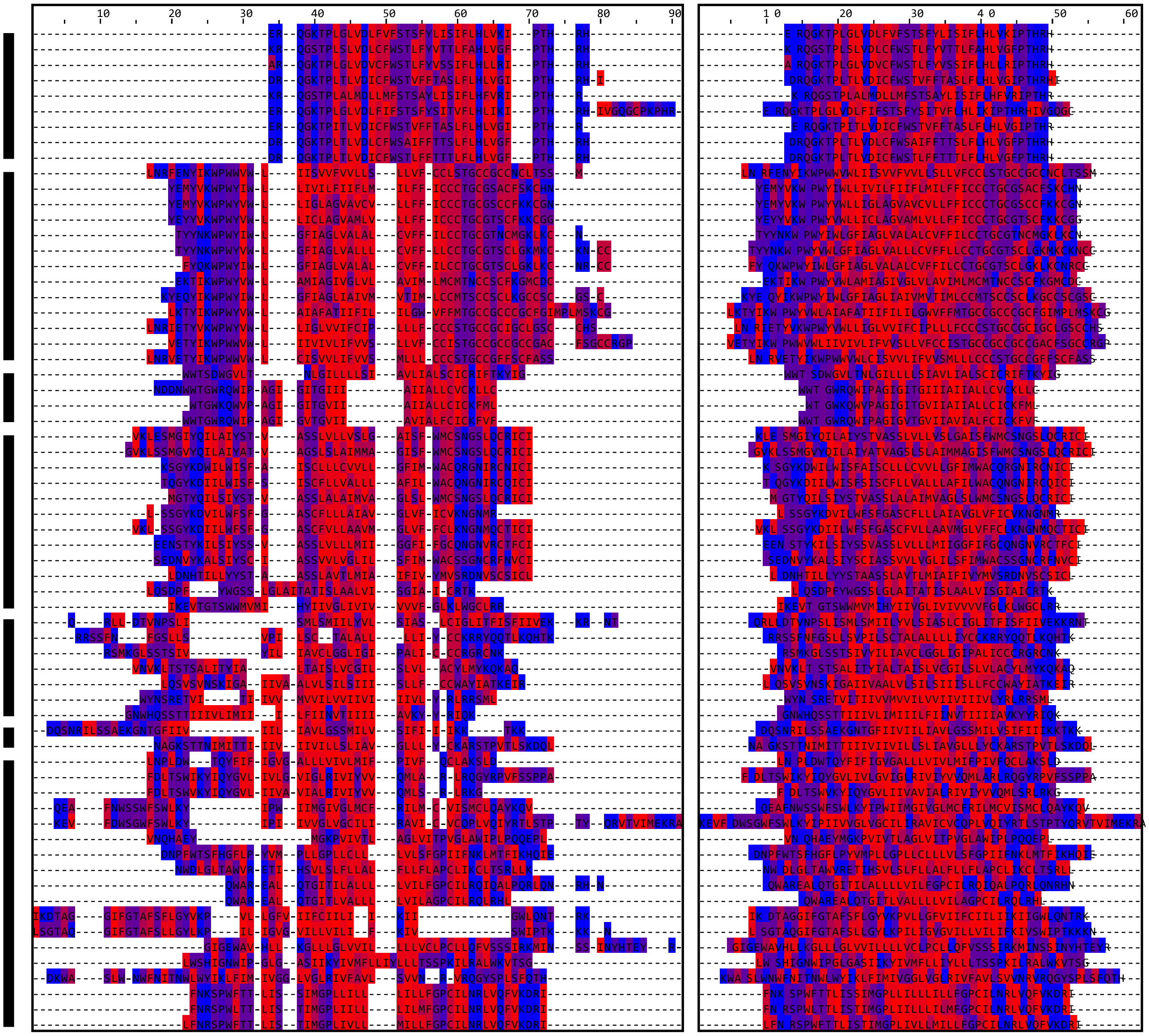

**Figure S5: Two different alignments of the predicted TM region of the cI-VFP dataset.** The predicted superset sequences from the cI-VFPs were padded on both sides by 5 residues and then aligned using Clustal omega (Sievers, et al., Mol Syst Biol, 2011; left panel) and manually at the center (right panel). The residues are colored using the hydrophobicity scale in Jalview (red: hydrophobic to blue: hydrophilic). The thick black bars on the left mark the different virus families (a-g) from top to bottom listed as in Table S1.

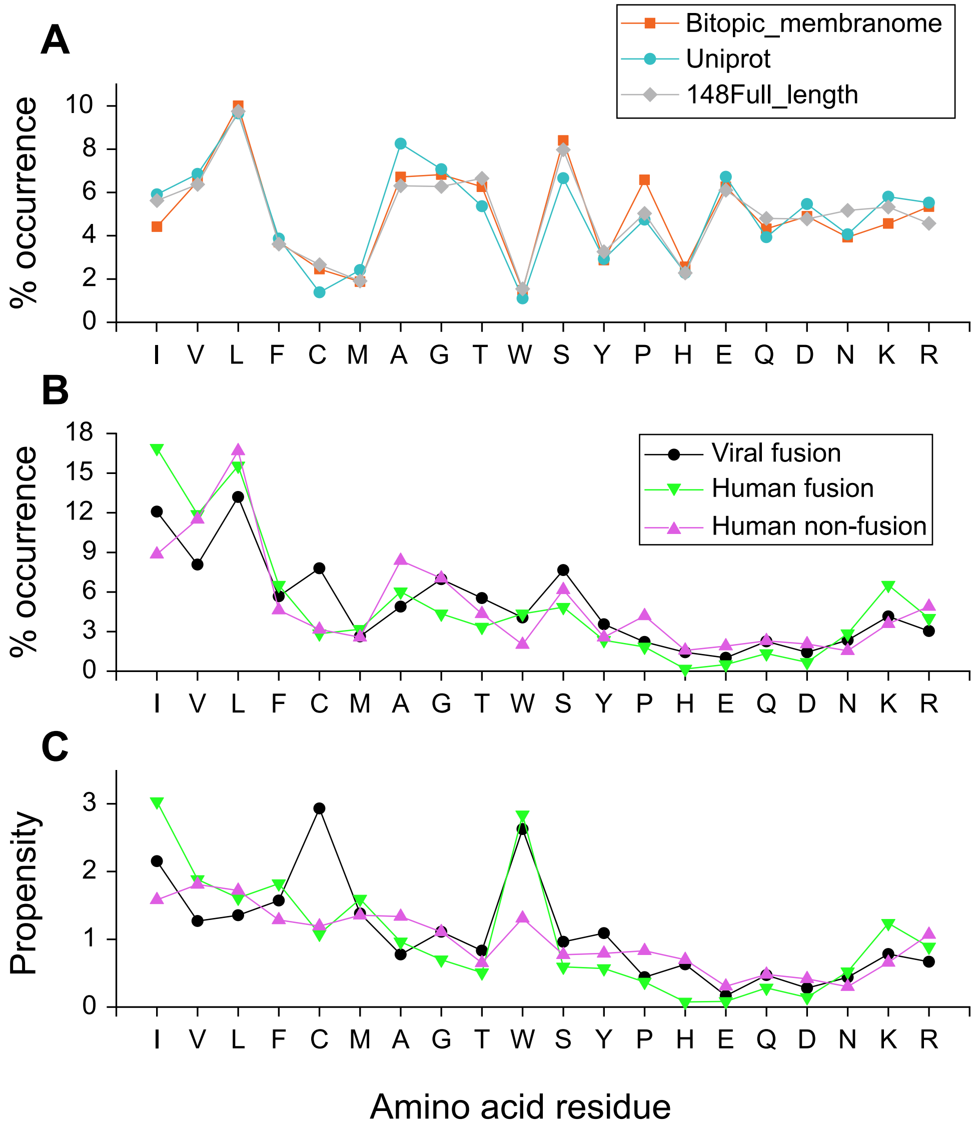


**Figure S6. Amino acid composition.** (**A**) The full-length amino acid composition of the spTMP membranome database (https://membranome.org/download/; updated 29th July, 2024), Uniprot statistics of the Reviewed Swiss-Prot database (https://www.uniprot.org/uniprotkb/statistics, as on 26th November, 2024), and the dataset of 148 spTMPs included in this study. The amino acid residues are arranged in the order of decreasing hydrophobicity according to the Kyte-Doolittle scale (Kyte and Doolittle, J Mol Biol, 1982) (**B**) The amino acid composition of the TM region (superset extended by five residues on both termini as in Fig. S5) for the three datasets used here: cl-VFPs (black), human fusion proteins (green), and human non-fusion proteins (pink). (**C**) The amino acid composition of the TM region in (B) normalized to the full-length amino acid composition of the 148 proteins in the dataset (gray diamonds in A).
